# Pharmacological profiling of a *Brugia malayi* muscarinic acetylcholine receptor as a putative antiparasitic target

**DOI:** 10.1101/2022.08.31.506057

**Authors:** Kendra J Gallo, Nicolas J Wheeler, Abdifatah M Elmi, Paul M Airs, Mostafa Zamanian

## Abstract

The diversification of anthelmintic targets and mechanisms of action will help ensure the sustainable control of nematode infections in response to the growing threat of drug resistance. G protein-coupled receptors (GPCRs) are established drug targets in human medicine but remain unexploited as anthelmintic substrates despite their important roles in nematode neuromuscular and physiological processes. Bottlenecks in exploring the druggability of parasitic nematode GPCRs include a limited helminth genetic toolkit and difficulties establishing functional heterologous expression. In an effort to address some of these challenges, we profile the function and pharmacology of muscarinic acetylcholine receptors in the human parasite *Brugia malayi,* an etiological agent of human lymphatic filariasis. While acetylcholine-gated ion channels are intensely studied as targets of existing anthelmintics, comparatively little is known about metabotropic receptor contributions to parasite cholinergic signaling. Using multivariate phenotypic assays in microfilariae and adults, we show that nicotinic and muscarinic compounds disparately affect parasite fitness traits. We identify a putative G protein-linked acetylcholine receptor (*Bma*-GAR-3) that is highly expressed across intra-mammalian life stages and adapt spatial RNA *in situ* hybridization to map receptor transcripts to critical parasite tissues. Tissue-specific expression of *Bma-gar-3* in *Caenorhabditis elegans* (body wall muscle, sensory neurons, and pharynx) enabled receptor deorphanization and pharmacological profiling in a nematode physiological context. Lastly, we developed an image-based feeding assay as a reporter of pharyngeal activity to facilitate GPCR screening in parasitized strains. We expect that these receptor characterization approaches and improved knowledge of GARs as putative drug targets will further advance the study of GPCR biology across medically important nematodes.

## Introduction

Parasitic nematodes cause infectious diseases of poverty endemic to underdeveloped and exploited countries, accounting for the loss of over 8 million disability-adjusted life years [1]. Current control mechanisms for helminth infections rely on mass drug administration (MDA) with a limited arsenal of drugs. Lymphatic filariasis (LF) is a neglected tropical disease caused by mosquito-transmitted nematodes *(Wuchereria bancrofti, Brugia malayi,* and *Brugia timori)* that migrate to and develop in human lymphatics [2]. An estimated 50 million people currently have LF with at least 36 million people suffering from chronic debilitating and highly stigmatizing conditions such as elephantiasis and hydrocele [3–6]. Anthelmintics used for LF treatment are suboptimal; they do not kill adult stage parasites and are contraindicated in regions co-endemic for closely related parasites. Further, the threat of anthelmintic resistance [7–13] underscores a recognized need for new drugs to treat vector and soil-transmitted nematode infections in human and animal populations.

Current anthelmintics were primarily discovered using animal or whole-organism screening approaches [14,15] and no new anthelmintics have been approved for human use in decades. Target-based approaches may provide an alternative route to screening validated molecular targets at much higher throughput [15–17], but bottlenecks derive from limited knowledge of basic parasite biology, a dearth of actionable targets, and difficulties in establishing reliable heterologous platforms for target expression and screening [16,18]. While ligand-gated ion channels (LGICs) receive warranted attention as the primary targets of existing anthelmintics, there is a need to diversify and pursue other druggable proteins critical to the physiology and survival of parasitic nematodes [9,19–22].

G protein-coupled receptors (GPCRs) are highly druggable and are the targets of over one-third of all FDA approved drugs in human medicine [23]. Despite their recognition as lucrative targets [24–29], helminth GPCRs have yet to be effectively exploited as anthelmintic substrates. Studies of GPCRs and their ligands in free-living nematodes show that this receptor family is involved in a range of important physiological processes [30–34]. Biogenic amines and neuropeptides elicit phenotypes of interest in free-living [30,31,35,36] and parasitic nematodes [26,37–39], many of which are likely mediated by metabotropic receptors. However, there is little data on the localization and function of parasitic nematode GPCRs, and pharmacological data is scant partly owing to difficulties in establishing reliable heterologous expression in single-cell systems [18,40,41]. Methods to characterize GPCRs in less tractable parasite species will better enable the prioritization of new receptor leads with host-divergent pharmacological profiles that can be selectively targeted.

Acetylcholine (ACh) and its receptor targets are essential for growth, development, and neuromuscular function in the clade V model nematode *Caenorhabditis elegans* [42,43]. The contribution of nicotinic acetylcholine receptors (nAChRs) to cholinergic signaling is underscored by the successful development of nicotinic channel agonists as antiparasitics [44–50], but much less is known about the druggability of muscarinic acetylcholine receptors (mAChRs) associated with slower but more sustained synaptic and extrasynaptic transmission.

The *C. elegans* genome encodes three known G protein-linked acetylcholine receptors (GARs) [51–54] that are widely expressed in the nervous system and muscle tissues [53,55] and are involved in the regulation of feeding, mating, egg laying, and locomotion [32,52,56–58]. While some of the GAR functions in *C. elegans* are likely conserved in parasitic nematodes, very little is known about GAR biology in filariae and other clade III parasites. GAR-1 from the gastrointestinal nematode *Ascaris suum* displays atypical pharmacologic responses [25,59] and muscarinic compounds affect motility in adult stage *Brugia malayi* [60], justifying closer examination of the GAR receptor subfamily.

Here, we focus our efforts on the characterization of a phylum-conserved muscarinic acetylcholine receptor in *Brugia malayi (Bma-gar-3).* We examine the effects of muscarinic compounds on microfilariae and adult *Brugia* using multivariate phenotyping approaches, and determine temporal and spatial gene expression patterns for *Bma-gar-3.* Building on previous work [35,61–64], we exploit the physiological context of *C. elegans* as a versatile heterologous platform for the study and characterization of parasite GPCRs as anthelmintic targets. We establish functional expression of *Bma*-GAR-3 in *C. elegans* and tissue-specific phenotypic endpoints in parasitized strains that allow for deorphanization and pharmacologic characterization. Lastly, we validate a high-throughput assay that enables screening of *Bma*-GAR-3 expressed in the *C. elegans* pharynx. These approaches circumvent some of the challenges associated with the study of GPCRs in difficult helminth systems, and are likely extensible to many other parasitic nematodes and receptors.

## Results and Discussion

### *Brugia malayi* GAR-3 is highly expressed across intra-mammalian life cycle stages and may mediate whole-organism effects of muscarinic compounds

Homology-based searches of annotated *C. elegans* G protein-linked acetylcholine receptors (GARs) were used to identify closely related biogenic amine receptors across six parasitic nematode species. Phylogenetic analysis of putative GARs reveals *B. malayi* possesses one-to-one orthologs of *C. elegans* GAR-2 and GAR-3 but not GAR-1 (**Figure 1A**). The clade IIIb nematode *Ascaris suum* possesses a GAR-1 ortholog, suggesting that GAR-1 was lost in the *B. malayi* clade IIIc sublineage. Although GAR-3 clusters closest to human mAChRs [65], nematode GARs are significantly diverged from their mammalian host orthologs and likely exhibit distinct pharmacological profiles that may allow for selective targeting [25,53,66]. In order to determine temporal patterns of *B. malayi* GAR expression, we performed qPCR across intra-mammalian life cycle stages (microfilariae (mf), L3, adult male, and adult female). *Bma-gar-3* is constitutively expressed across all life cycle stages and is much more highly expressed than *Bma-gar-2* (**Figure 1B**), suggesting potentially outsized physiological roles throughout development.

**Figure 1.**
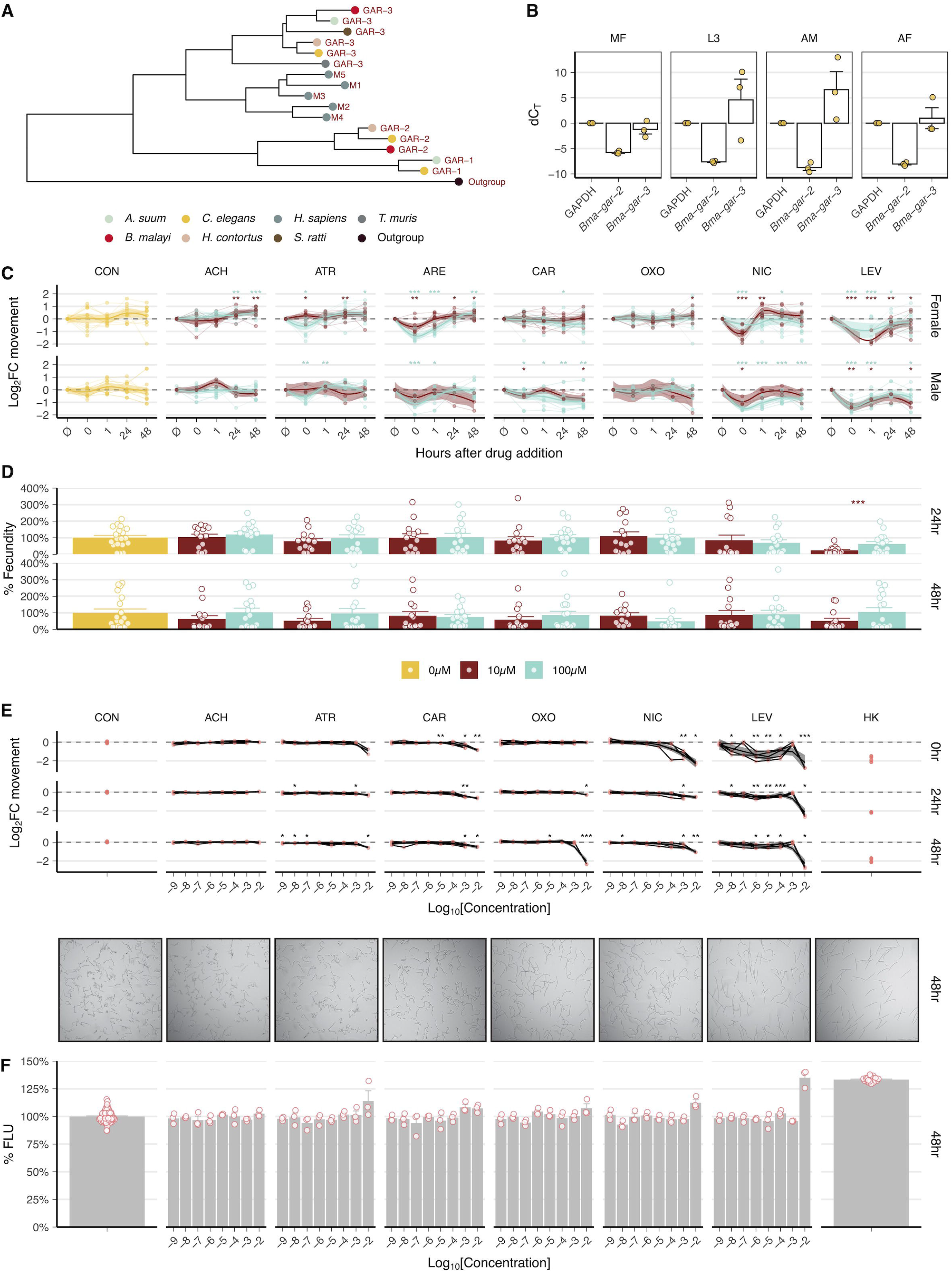
*B. malayi* expresses two GARs and muscarinic compounds elicit neuromuscular effects in microfilariae and adult stage parasites. **A)** Phylogeny based on protein sequence alignment of characterized and putative nematode and human GARs. *B. malayi* expresses two GARs (*Bma*-GAR-2 and *Bma*-GAR-3), but lacks a GAR-1 homolog. **B)** Expression of *Bma-gar-2* and *Bma-gar-3* across intra-mammalian life stages of *B. malayi* as assayed by quantitative reverse transcriptase PCR (qPCR) (control = *B. malayi* GAPDH). *Bma-gar-3* is abundantly and constitutively expressed throughout the life cycle. **C)** Effects of cholinergic compounds on adult *B. pahangi* movement over 48 hours as measured by optical flow. Log2 normalized movement compared to untreated controls using a t-tests (*p ≤ 0.05; **p ≤ 0.01, ***p ≤ 0.001) (brown: 10 μM, cyan: 100 μM). **D)** Effects of cholinergic compounds on adult female *B. pahangi* fecundity as measured by microfilariae output. Percent change mf output from untreated controls by time point, t-tests (*p ≤ 0.05; **p ≤ 0.01, ***p ≤ 0.001) (brown: 10 μM, cyan: 100 μM). **E)** Dose-response of cholinergic compounds on microfilariae motility over 48 hours. Images from mf motility plates taken at 48 hours showing diverse morphologies from cholinergic treatment at 10 mM. Log2 normalized movement compared to untreated controls using t-tests (*p ≤ 0.05; **p ≤ 0.01, ***p ≤ 0.001). **F)** Effects of cholinergic compounds on microfilariae cell health over 48 hours as measured by cell toxicity stain (CellTox fluorescence). Percent change in fluorescence from untreated controls, t-tests (*p ≤ 0.05; **p ≤ 0.01, ***p ≤ 0.001).

To explore the gross effects of cholinergic compounds on parasite health, we optimized a number of phenotypic readouts in *Brugia* adults and microfilariae [67]. Parasites were incubated in muscarinic and nicotinic compounds and the stage-specific effects of these chemical perturbations on worm motility, viability, and fecundity were measured using a customized imaging platform [68]. Male and female *Brugia* adults were exposed to 10 μM and 100 μM of acetylcholine (ACh), atropine (ATR), arecholine (ARE), carbachol (CAR), oxotremorine-M (OXO), nicotine (NIC), and levamisole (LEV). ACh leads to a slow increase in baseline movement after prolonged exposure (24 and 48 hours), attributable to inefficient penetration of the lipid cuticle [69] (**Figure 1C**). Treatment with nicotinic compounds (NIC and LEV) leads to an immediate drop in female and male (10 μM and 100 μM) worm motility followed by quick recovery, mediated by fast responding nAChRs [70].

Treatment with compounds associated with muscarinic activity (ATR, ARE, CAR, OXO) elicit a range of subtle-to-large effects on motility. ATR and ARE immediately decrease motility in male and female worms (100 μM) and CAR decreases motility in male worms (10 μM and 100 μM), in agreement with previous work [60]. *Cel-*GAR-2 is unaffected by the muscarinic agonists ARE and OXO or the antagonist ATR [53,65], suggesting that effects driven by these compounds are mediated by *Bma*-GAR-3. Given the promiscuity of some muscarinic compounds, it is possible that some of these effects are partly mediated by nicotinic receptors or that acute effects are dominated by ionotropic as opposed to metabotropic signaling. Fecundity was not significantly altered by any of the treatments except for 10 μM LEV at 24 hrs, a nicotinic compound that stimulates egg-laying in *C. elegans* [71] (**Figure 1D**).

Dose responses were carried out in mf stage parasites over three time points (0, 24, and 48 hours) to measure the effects of cholinergic compounds on motility and cell death. NIC and LEV both cause immediate inhibition of mf motility at high (>10^-4^ M) and low (>10^-7^ M) concentrations, respectively (**Figure 1E**). OXO, a GAR-3 selective muscarinic compound, significantly decreases motility at high concentrations (>10^-2^ M) in the 24-48 hour time frame. Morphologies of mf at 48 hours varied among treatments suggesting effects not fully captured by motility. LEV, NIC, and OXO treatment causes worms to become flaccid much like heat-killed (HK) control worms while ACh-treated worms maintain the posture of untreated controls. Some cell death was noticeable for all treatments except acetylcholine at high concentrations (>1 mM) at 48 hours (**Figure 1F**). While OXO-mediated effects suggest that perturbation of *Bma*-GAR-3 may elicit phenotypes of interest, the slower neuromodulatory action of this general receptor class may require assays sensitized to other subtle but important phenotypes in the host context [72,73]. More insight into tissue-specific expression patterns of this highly-expressed receptor would allow for better prediction of its physiological roles.

### *Bma-gar-3* transcripts are widely expressed across critical tissues in adult stage parasites

While GARs have been localized to specific cells and tissues using genetic tools in the tractable *C. elegans* system, the fine spatial distribution of these receptors is unknown in filarial or other parasitic nematodes. Building on an RNA tomography protocol [74], we developed a strategy to map *Bma-gar-3* transcripts across the adult female head and mid-section at 8 μm resolution while preserving spatial information (https://doi.org/10.6084/m9.figshare.20757481.v1). Individual sections were sequentially captured from the anterior tip (216 sections) of a single adult female *B. malayi* worm and RNAscope was used to localize *Bma-gar-3* transcripts within sections, allowing for reconstruction of expression patterns down the anterior-posterior axis of the head region (**Figure 2A**). *Bma-gar-3* is widely expressed across several tissue types including the body wall muscle and neurons, with near ubiquitous expression in digestive and reproductive tissues (**Figure 2B-D**). The expression of *Bma-gar-3* across these important tissues overlaps with the expression pattern of *C. elegans gar-3* (body wall muscle, pharyngeal muscle, cord and other neurons) [75–78], suggesting some conservation of pleiotropic receptor function across clades.

**Figure 2.**
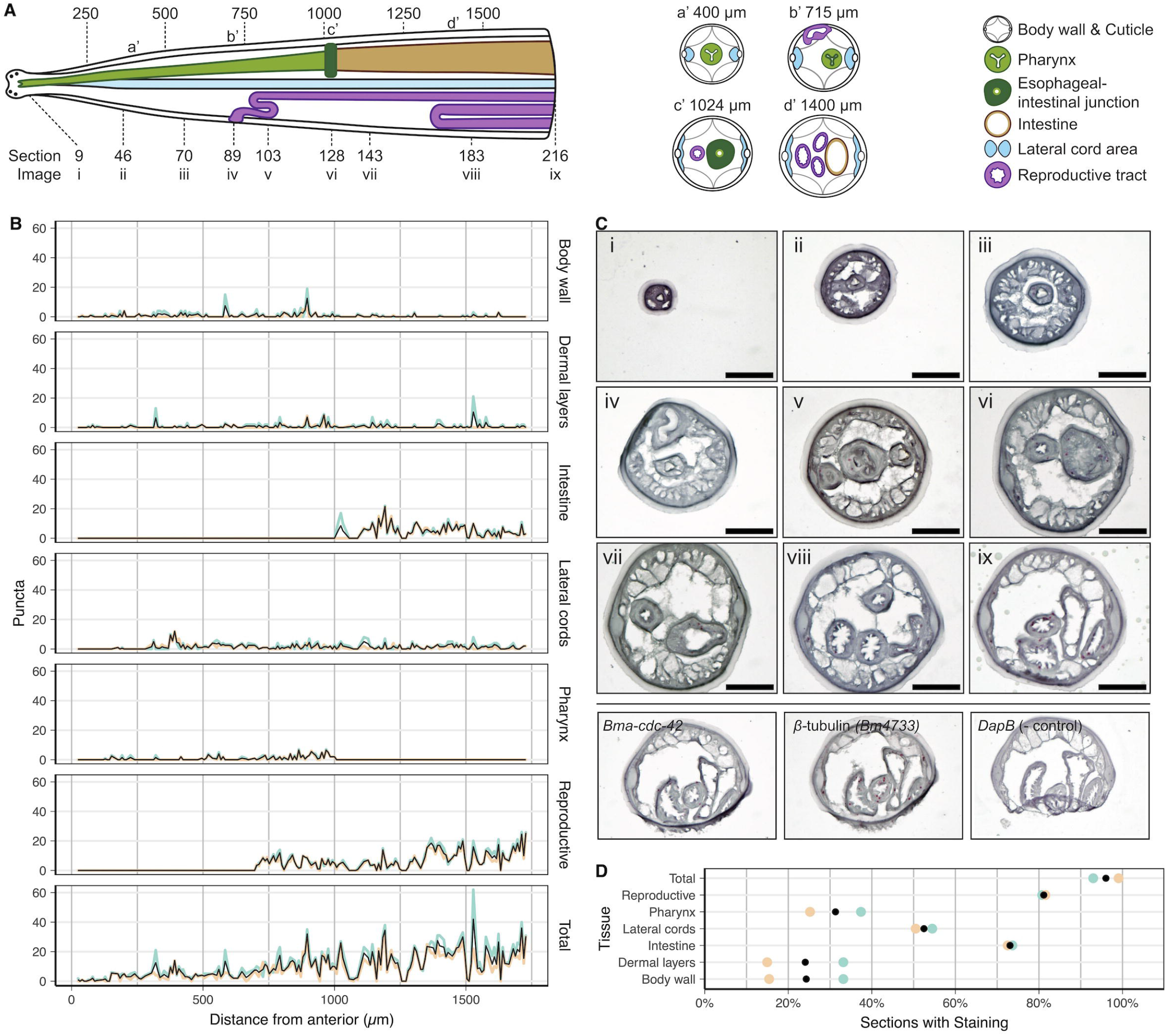
RNAscope spatial localization of *Bma-gar-3* in the *B. malayi* adult female head. **A)** Illustration of tissue distribution along the anterior-posterior axis and transverse illustrations at approximately (a’) 400 μm, (b’) 715 μm, (c’) 1024 μm, and (d’) 1400 μm, with a key of representative section locations shown in part C. **B)** RNAscope *Bma-gar-3* punctae counts per tissue per 8 μm section. **C**) Representative section images with key in part A, showing 3x zoom insets of punctate staining in the pharynx (iii), vulva (iv), uterus (v), esophageal-intestinal junction (vi), body wall muscle (vii), intestine (viii), and lateral cords (ix). Scale bar = 50 μm. Positive *(Bma-cdc-42* and *Bm4733)* and negative (bacterial *DapB)* controls for RNAscope validate RNA integrity, show that punctate staining is probe specific, and that nonspecific background staining is minimal. **D)** Proportion of sections containing *Bma-gar-3* punctae where tissue is present and defined. Colors in C and D represent counts performed by two independent researchers.

### Heterologous expression and deorphanization of *Bma-GAR-3* in *C. elegans*

We sought to establish heterologous assays to characterize the pharmacology of *Brugia malayi* GPCRs through functional expression in discrete *C. elegans* tissues. Pharmacological profiling of parasite GPCRs in this heterologous system requires that the receptors are properly folded and exported to the membrane, that they signal through endogenous G proteins, and that their activation in response to exogenous ligands can be measured through convenient phenotypic endpoints. Building on previous work leveraging *C. elegans* as a heterologous expression platform for studying anthelmintic targets [35,61–63], we first established transgenic lines expressing *Bma-*GAR-3 in the *C. elegans* ASH sensory amphid neuron and the body wall muscle. These parasitized strains were used to develop and optimize tissue-specific assays to measure receptor activation.

To limit background signaling, transgenic lines were created in a *Cel-*GAR-3 knockout *(gar-3(gk305))* genetic background. We employed simple plate-based assays to verify proper cell surface expression of *B. malayi* GAR-3 and to deorphanize the receptor by confirming activation by the putative ligand ACh. Activation of the *C. elegans* ASH neuron by noxious stimuli results in a well-characterized avoidance response whereby worms reverse their movement. We hypothesized that the successful activation of parasite GPCRs expressed in this neuron should lead to increased reversal frequency. We adapted an aversion assay [79,80] that involved placing individual worms in the center of a compound ring and monitoring for reversals in movement in response to test compounds **(Figure 3A)**. Worms expressing *Bma*-GAR-3 in the ASH neuron *(sra-6p::Bma-gar-3)* exhibited strong aversion responses to ACh (100 mM) and the selective muscarinic agonist oxotremorine-M (100 mM) **(Figure 3B)**. OXO has been shown to specifically activate *C. elegans* GAR-3, but not *Cel*-GAR-1 or *Cel-*GAR-2 [51,53,66]. Neither the wild-type (N2) or knock-out *(gar-3(gk305))* strains demonstrated aversion to ACh or OXO but all strains maintained consistent responses to negative (water) and positive (4 M fructose) controls.

**Figure 3.**
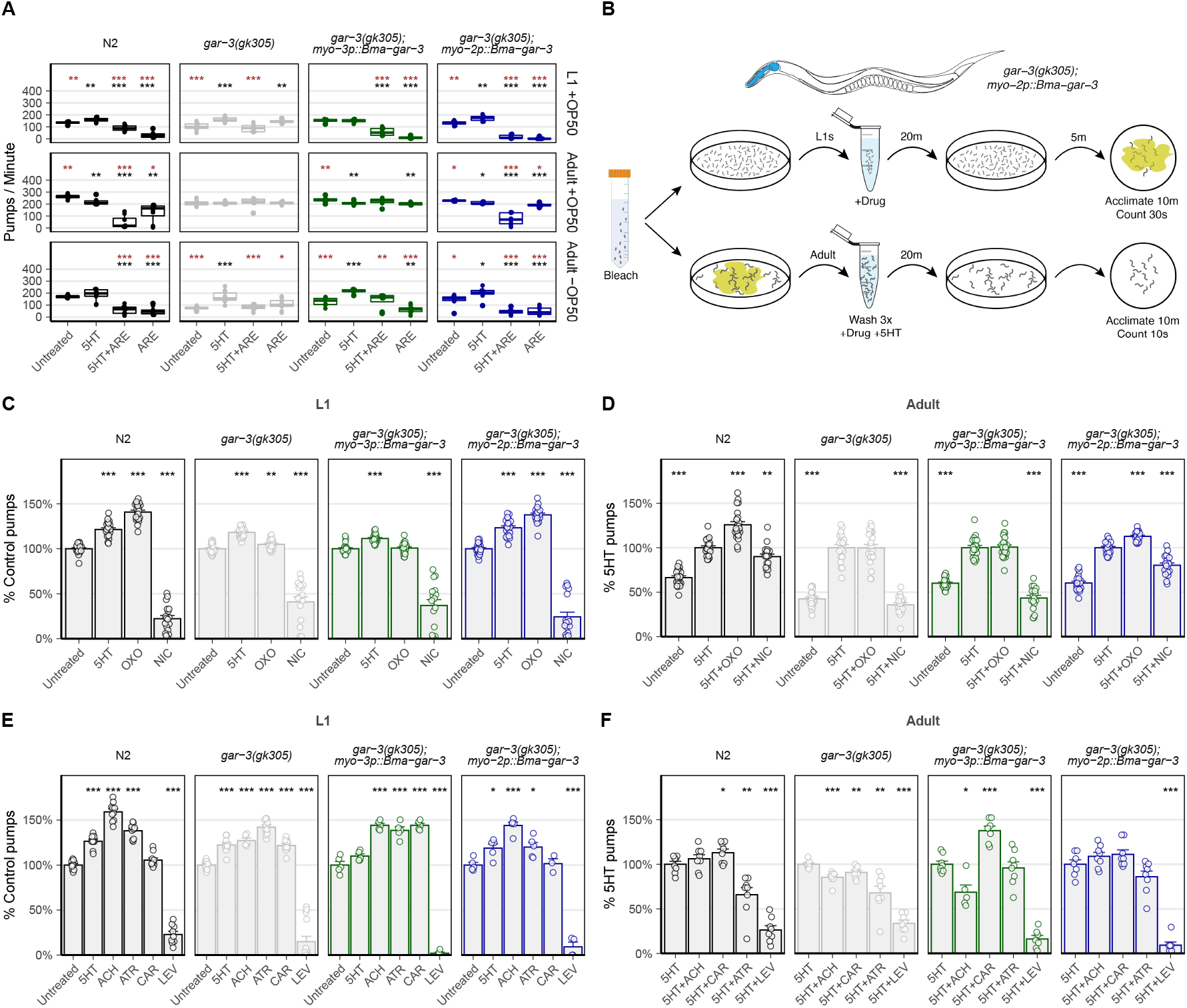
*Bma-GAR-3* is activated by the selective muscarinic agonist oxotremorine-M and confers hypersensitivity to aldicarb induced paralysis. A) Schematic of plate-based aversion assay. Bleach-synchronized adult worms are monitored to capture reversal frequency in response to a test compound ring, in the presence of a known attractant (diacetyl). **B)** Acetylcholine and oxotremorine-M activate the ASH neuron of parasitized *C. elegans* expressing *Bma*-GAR-3 and elicits reversal behaviors. t-tests (*p ≤ 0.05; **p ≤ 0.01, ***p ≤ 0.001) were used to identify differences in comparison to *gar-3(gk305)* (gray) and N2 (black). **C)** Schematic of aldicarb paralysis assay. Paralysis of young adult worms is monitored on agar plates containing test drug over a two hour period. **D)** Kaplan-Meier survival plot showing paralysis measured every 10 minutes. Knock-out of *Cel*-GAR-3 leads to resistance to aldicarb induced paralysis. *myo-3p::Bma-gar-3* worms paralyze more rapidly than wild-type N2 and *gar-3(gk305)* worms. Pairwise survival t-tests (*p ≤ 0.05; **p ≤ 0.01, ***p ≤ 0.001).

We then assayed the effects of a cholinesterase inhibitor (aldicarb) on worms expressing *Bma*-GAR-3 in the body wall muscle *(myo-3p::Bma-gar-3),* predicting that the build up of ACh at the receptor synapse would lead to flaccid paralysis of this parasitized *C. elegans* strain. We adapted a protocol [81] that involved transferring worms onto plates with 1 mM aldicarb, restricting their movement with copper rings, and monitoring responses to touch stimuli over a 120 minute period. *myo-3p::Bma-gar-3* worms are hypersensitive to aldicarb-induced paralysis compared to wild-type and knock-out strains **(Figure 3C-D)**. Combined, these results show that *Bma*-GAR-3 can be functionally expressed in both sensory neurons and body wall muscle, and that this receptor is activated by acetylcholine and oxotremorine-M.

### Establishing pharyngeal endpoints for profiling *Bma-GAR-3* pharmacology

In order to improve the pharmacologic profiling of parasite GPCRs in this heterologous system, we optimized more quantitative assays to measure receptor activation in response to exogenous drugs. Any eventual anthelmintic screen of parasite receptors expressed in this model system will require quantitative and scalable phenotypic readouts of receptor activity. We generated a strain of *C. elegans* expressing *Bma*-GAR-3 in the pharyngeal muscle *(myo-2p::Bma-gar-3),* hypothesizing that expression and activation of parasite acetylcholine receptors in the pharynx and the body wall would alter baseline and drug-induced pharyngeal pumping activity. Pharyngeal pumping activity can be directly measured by observing terminal bulb inversions [56,75] or using electropharyngeogram (EPG) recordings [82,83]. Each pump is highly regulated [57,84] by muscarinic receptors working in tandem with nAChRs [57,75,85].

To optimize assays that rely on pharyngeal function as a quantitative measure of direct or indirect parasite receptor activity, we investigated the effects of pumping stimuli across worm developmental stages on our ability to resolve drug responses in parasitized strains. We first examined how different pharyngeal pumping stimuli would affect our ability to measure drug-induced changes in both larvae and adults. While OP50 and serotonin (5HT) are commonly used to elevate baseline pumping frequency [82,86–90] for assay sensitization, it is important that these stimuli do not mask drug effects and allow for the reliable capture of both inhibitory and stimulatory responses to drug exposure. We tested combinations of OP50, 5HT, and a known GAR-3-dependent inhibitor of pharyngeal pumping (arecoline) [75] to assess our ability to capture effects in both directions in L1 and young adult animals.

L1 assays carried out on OP50 plates confirmed that this food stimulant does not saturate pumping responses or mask expected drug effects. The inhibitory effect of arecoline can be measured in strains expressing either *C. elegans* GAR-3 (N2) or *B. malayi* GAR-3 *(myo-2p::Bma-gar-3)* in the pharynx (**Figure 4A**). A dynamic range of pump frequencies is observable using OP50 and we conclude from these data that L1 assays should be carried out in the presence of OP50 and without 5HT. OP50 leads to a non-saturating increase in L1 baseline pharyngeal pumping that allows us to measure both drug-induced stimulation and inhibition of pump frequency. In contrast, adult stage assays carried out in the presence of OP50 saturated baseline pharyngeal pumping frequency. The addition of 5HT leads to no further increase in pumping rate and the inhibitory effects of ARE are largely masked in these conditions (**Figure 4A**). Assays carried out in the absence of OP50 allow for the robust detection of both 5HT stimulation and arecoline inhibition. We conclude from these data that adult assays should be carried out in the absence of OP50 and with drugs in combination with 5HT to allow for the largest dynamic range of stimulation and inhibition.

**Figure 4.**
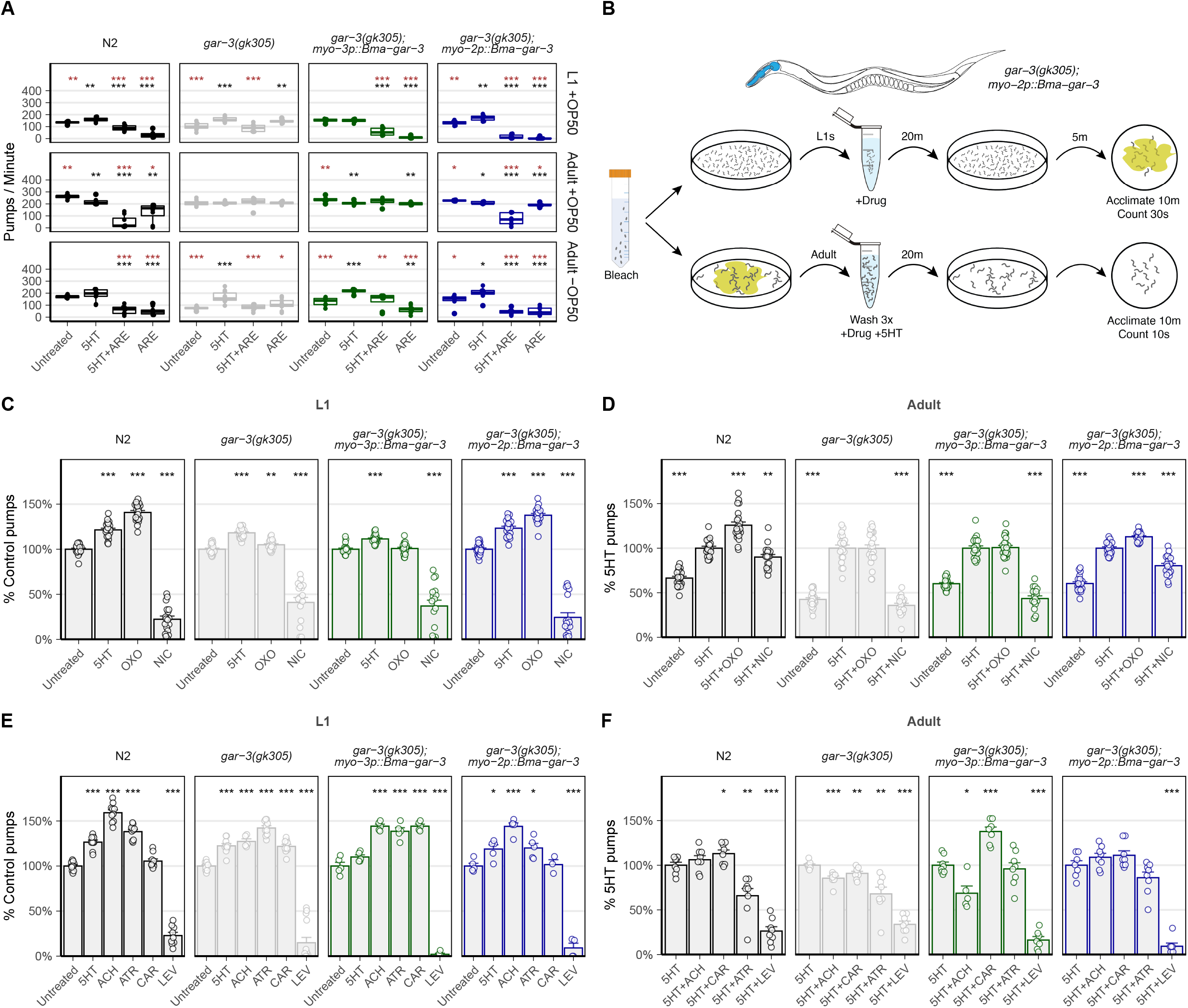
*Bma*-GAR-3 modulates pharyngeal pumping via expression in the *C. elegans* body wall and pharynx. **A)** Iteration across chemical (5HT) and food (OP50) pumping stimuli for optimization of visual pumping assay in L1 and adult stage *C. elegans.* (brown: comparisons to 5HT, black: comparisons to untreated). **B)** Schematic of the optimized visual pharyngeal pumping assay. Bleach synchronized worms are treated with drug for 20 minutes and pumps are counted on seeded (L1s) or unseeded (adults) 6 cm agar plates. Drug treatments are tested in combination with 5HT in the adult stage. **C-D)** Modulation of pharyngeal pumping by OXO in L1 and adult parasitized strains. L1 data are normalized to untreated controls while adult data are normalized to 5HT-stimulated controls. **E-F)** Effects of cholinergic compounds on stage specific pharyngeal pump frequency compared to untreated (L1) or 5HT-stimulated (adult) pump frequency. All statistics were calculated using t-tests (*p ≤ 0.05; **p ≤ 0.01, ***p ≤ 0.001).

We next used these optimized L1 and adult stage assays (**Figure 4B**) to confirm the action of the selective *Cel-*GAR-3 agonist oxotremorine-M. OXO increases pumping frequency in L1 (~ 41%) and adult (~26%) stage N2 worms (**Figure 4C-D**). Knock-out of native *gar-3* leads to a loss of OXO responsiveness, which is near completely rescued by expression of *Bma-gar-3* in the pharynx but not the body wall muscle. We next profiled the responses of parasitized strains to muscarinic and nicotinic compounds with less receptor specificity. In L1 stage worms, expression of *Bma-gar-3* in the pharynx restores the wild-type response profile (**Figure 4E**). In adult stage worms, expression of *Bma-gar-3* in the body wall leads to hyperstimulation of pumping in response to CAR, while expression of *Bma-gar-3* in both the pharynx and body wall leads to a decreased inhibitory response to ATR compared with either wild-type or *gar-3(gk305)* strains (**Figure 4F**).

Although it is known that pharyngeal pumping can be modulated by cholinergic signaling in both tissue types, the precise mechanism by which the body wall and pharynx communicate is unclear [83,91,92]. Interpretations of how pharyngeal pumping is modulated by direct versus indirect pharmacological action at our receptor of interest can be confounded by the promiscuous binding of cholinergic compounds to a range of muscarinic and nicotinic receptors expressed across relevant tissues and perhaps differentially expressed across stages. Despite these complications, it is reasonable to expect that compounds with high specificity for GAR-3 can be identified by comparing responses across *gar-3(gk305)* and *gar-3(gk305); myo-2p::Bma-gar-3* strains.

### Electropharyngeal measurements of *Bma*-GAR-3 activity in parasitized strains

Electrophysiological recordings from the pharynx can provide more detailed information about pharyngeal function in response to drug. Using established protocols [82], we sought to use electropharyngeogram (EPG) recordings to investigate other pharyngeal phenotypes modulated by *Bma-*GAR-3 expression in the body wall and pharynx of young adult *C. elegans.* We recorded individual worms for two minutes after a 20 minute drug incubation period, mirroring the visual counting assay (**Figure 5A**). The use of 5HT in combination with test drugs was necessary to capture the inhibitory effects of ARE and to recapitulate trends from visual counting data (**Figure 5B**).

**Figure 5.**
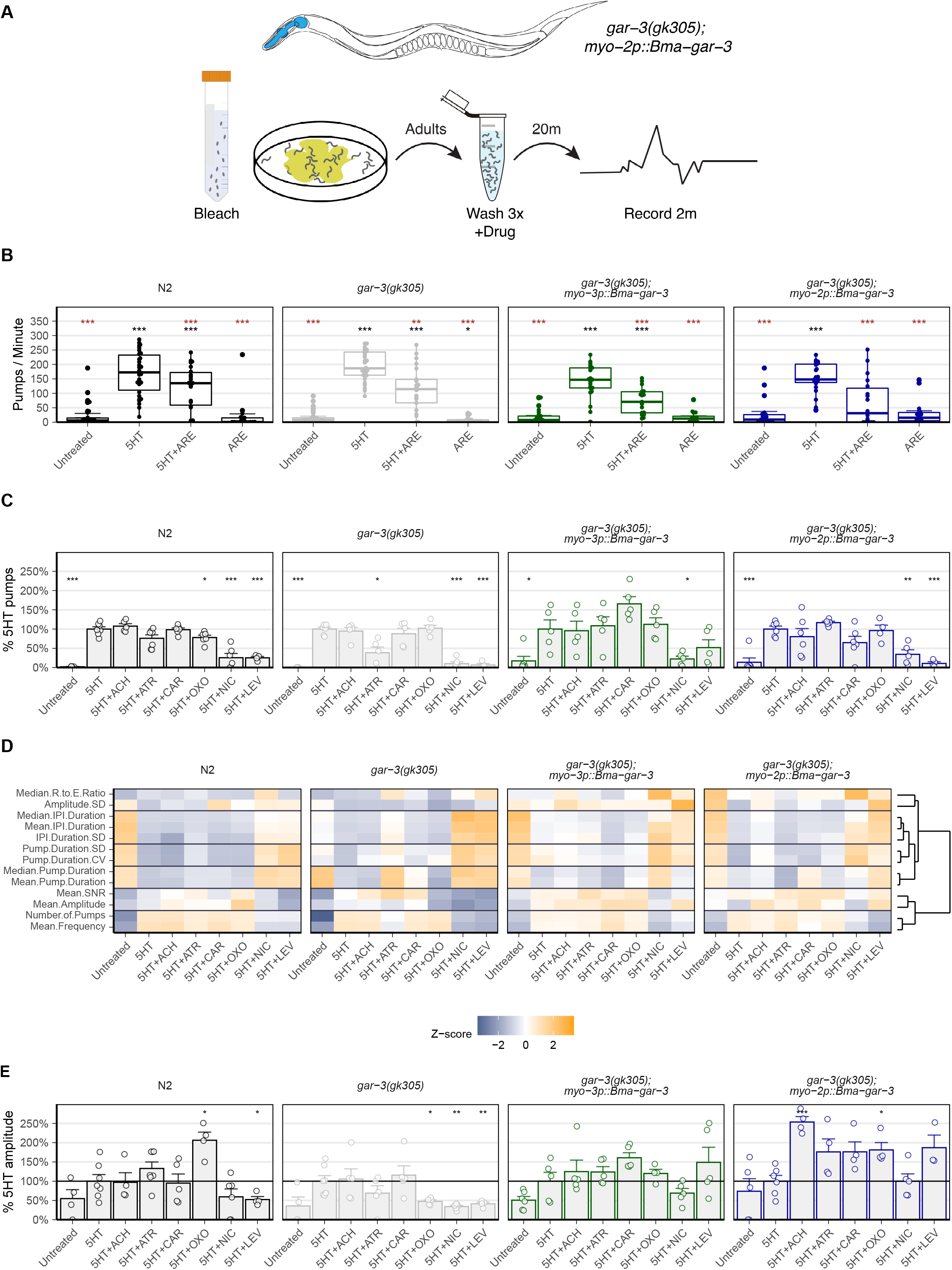
EPG recordings in parasitized strains provide alternative reporters of parasite receptor activity. **A)** Schematic of electropharyngeogram (EPG) assay. Adult worms are bleach synchronized, washed three times, treated with drugs for 20 minutes, and positioned into the microfluidics chip. Worms are recorded for two minutes. **B)** EPG recordings used to measure pharyngeal pump frequency show that 5HT in combination with drugs allows for the capture of inhibitory effects of ARE. (brown: comparisons to 5HT, black: comparisons to untreated). **C)** Cholinergic effects on pharyngeal pump frequency as measured by EPG do not align with visual pumping assay. **D)** Heatmap depicting scale-normalized electrophysiological features across all strain conditions. Clustering of these features identifies subsets that provide similar information. **E)** *Bma*-GAR-3 expressed in the pharynx increases the peak amplitude of OXO treated worms compared to *gar-3(gk305)* knockout, revealing receptor-specific modulation of this electrophysiological feature. All statistics were calculated using t-tests (*p ≤ 0.05; **p ≤ 0.01, ***p ≤ 0.001).

We next used EPG recordings to test whether the effects of cholinergics on electrophysiological features could be linked to *Bma*-GAR-3 activity in parasitized strains. We found that EPG-derived pump frequency did not correlate well with visual counting data. Most notably, OXO did not exhibit a pattern of differential response and rescue in *gar-3(gk305)* and *gar-3(gk305);myo-2p::Bma-gar-3,* respectively (**Figure 5C**). General discrepancies between visual counts and EPG-derived pump frequency are likely due to the disconnect between terminal bulb movement and action potentials, supported by the fact that *C. elegans* GAR-3 regulates both membrane potential and excitation-contraction coupling through a yet defined signaling pathway [75]. While EPG-derived pump frequency was not a reporter of *Bma-*GAR-3 activation in this assay, other electrophysiologic features show patterns consistent with *Bma-*GAR-3 phenotypic rescue (**Figure 5D**). Specifically, expression of *Bma-gar-3* in the pharynx rescues the OXO-induced increase in peak amplitude that is lost in the *gar-3(gk305)* background (**Figure 5E**). EPG recordings provide a rich set of features (**Sup Figure 1**) that can reveal the activation of parasite receptors in response to exogenous drug. While these electrophysiological assays provide deeper insight into electric and chemical signaling dynamics, they do not enable high-throughput screening of parasitized strains to identify drugs that act on receptors of interest.

### Establishing a high throughput image-based feeding assay for screening of parasitized strains

We developed a high-throughput imaging assay to measure pharyngeal pumping as a reporter of receptor activity in parasitized animals. Fluorescence uptake in the form of beads, bacteria, and dye has been used to measure feeding behaviors in *C. elegans,* whereby drug modulation of pumping rates can be expected to impact the amount of intestinal fluorescence. Many of these assays are low in throughput [93–96] or require a large particle sorter [97–100], necessitating the development of a high-throughput and high-content imaging endpoint that allows for the measurement of intestinal fluorescence and transgenic markers.

We optimized parameters for a microtiter plate assay that measures intestinal accumulation of fluorescent dye as a correlate of pharyngeal activity. L1 synchronized worms were aliquoted into 96-well plates and grown to adults over 48 hours in the presence of HB101. Worms are then treated with a test compound for 20 minutes followed by a 20 minute BODIPY 558/568 (red) incubation. We tested two concentrations of BODIPY to help minimize background fluorescence and ensure that dye was not a limiting reagent. We tested the inclusion of 5HT and HB101 as feeding stimulants for the duration of the drug exposure, as well as the inclusion of HB101 in the BODIPY incubation period. Assays using at least 90 ng/mL BODIPY (2x) generated the best conditions to detect both pharyngeal stimulation (5HT and HB101) and inhibition (NIC) (**Figure 6A)**. The inclusion of HB101 in the dye incubation period was not necessary to detect these differences (**Figure 6A**).

**Figure 6.**
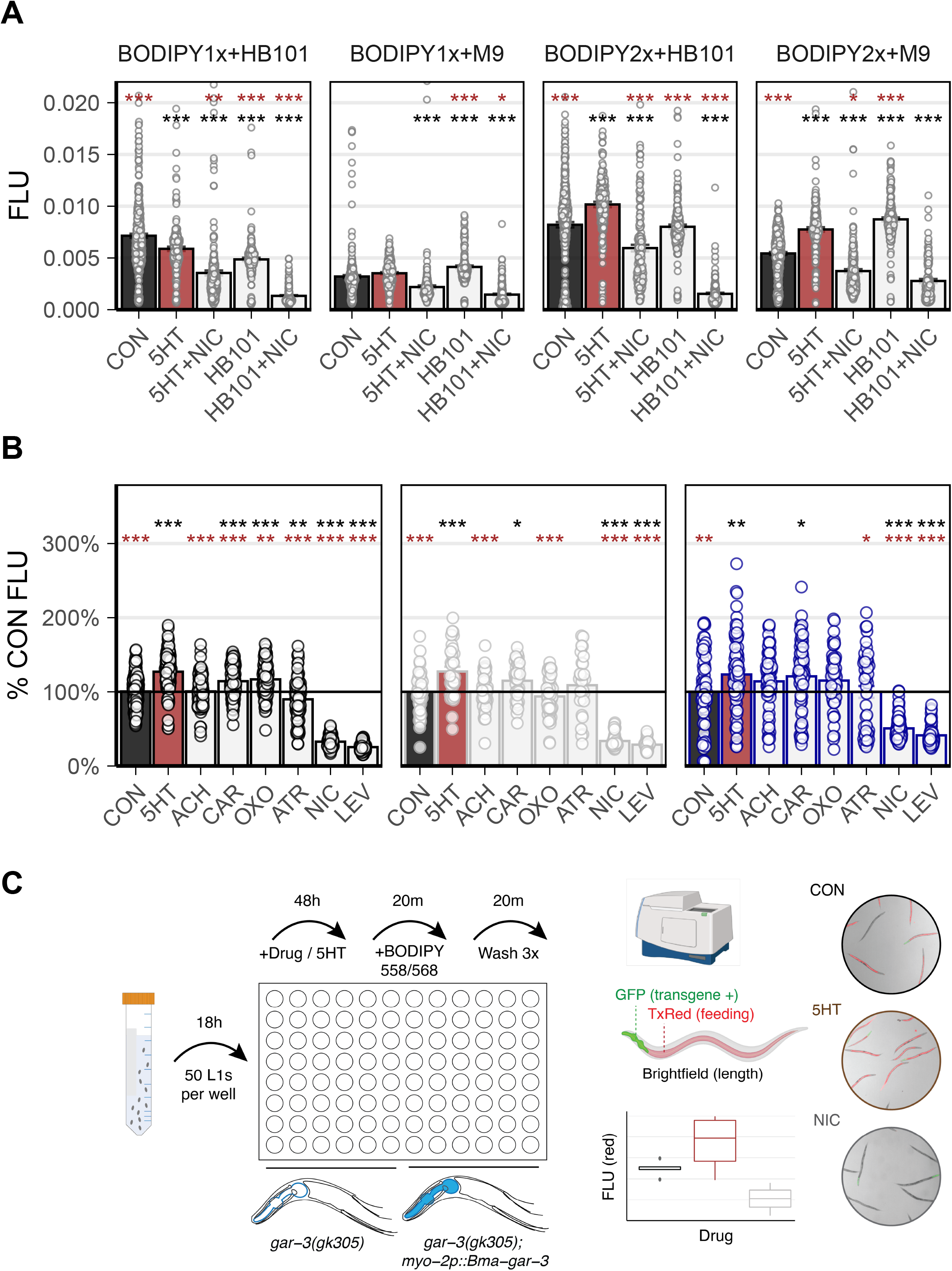
Development of an image-based feeding assay to enable high-throughput screening of parasitized strains. **A)** Testing lipophilic dye (BODIPY) concentration (1x: 45 ng/mL, 2x: 90 ng/mL) and the inclusion of HB101 for the development of a feeding assay. Treatment with 2x BODIPY in the absence of HB101 allowed the detection of both pharyngeal stimulation and inhibition as quantified by the accumulation of red intestinal fluorescence. (brown: comparisons to 5HT, black: comparisons to untreated). t-tests (*p ≤ 0.05; **p ≤ 0.01, ***p ≤ 0.001). **B)** Feeding assay carried out using a panel of cholinergic treatments. *Bma*-GAR-3 expression in the pharynx partially rescues the wild-type OXO effect that is lost in the *gar-3(gk305)* background, as measured by fluorescent dye uptake. (brown: comparisons to 5HT, black: comparisons to untreated). t-tests (*p ≤ 0.05; **p ≤ 0.01, ***p ≤ 0.001). C) Schematic of the dye feeding assay. Bleach synchronized L1 worms are grown in 96-well plates for 48 hours followed by 20 minute drug treatment. Worms are then fed >90 ng/mL BODIPY 558/568 for 20 minutes. Plates are washed three times and worms are paralyzed and straightened with 50 mM sodium azide before images are acquired using an high-content imaging system. Fluorescence is quantified using a wrmXpress[68] pipeline.

To validate this protocol as a means to screen parasite receptors at higher throughput, we compared the effects of cholinergic compounds in *gar-3(gk305)* and *gar-3(gk305); myo-2p::Bma-gar-3* animals. We treated worms with cholinergic compounds followed by BODIPY incubation. An image processing pipeline was established to identify transgene (+) worms via pharyngeal GFP expression. OXO causes an increase in dye uptake in wild-type worms compared to non-treated controls (~ 17%, p = 9.4e-07), consistent with increased pumping frequency observed in visual counting assays. OXO decreases dye uptake in the *gar-3(gk305)* background, which is rescued by expression of *Bma-gar-3* in the pharynx (**Figure 6B**). These results reaffirm that OXO has selective effects on *Bma-*GAR-3 in the pharynx and provides proof of principle for this high content imaging approach for pharmacological profiling of transgenically expressed parasite GPCRs.

## Conclusion

Parasite G protein-coupled receptors remain unexploited as anthelmintic targets despite their involvement in critical nematode neuromuscular and physiological processes. One significant bottleneck in exploring the pharmacology of parasite GPCRs results from difficulties in consistently establishing heterologous expression in single-cell systems. Yeast and mammalian cell culture systems have paved the way for deorphanization of helminth GPCRs [18,25,101], but not all receptors express or behave properly in cell types derived from distant phylogenetic lineages. The combinations of accessory proteins, molecular chaperones, G proteins, and membrane determinants required for the successful folding, cell-surface expression, and signaling of parasite receptors in surrogate systems have not been comprehensively identified. To avoid some of these complications, we explored a range of assays for parasite GPCR expression and profiling in the model nematode *C. elegans.*

We identified a *B. malayi* muscarinic GPCR *(Bma-gar-3)* that is highly expressed throughout the intra-mammalian life stages and developed a spatial RNAscope protocol to map receptor transcripts in multiple tissue types in the adult stage. Multivariate phenotypic assays of cholinergic effects on microfilariae and adult parasites reveal differential effects of nicotinic and muscarinic agents. We show that muscarinic compounds affect motility in both adult and mf stage parasites, some of which are likely to be mediated by GAR-3. We predict that sampling a broader array of muscarinic compounds will likely reveal other overt and subtle but important phenotypes that are relevant to potential anthelmintic mechanisms of action.

Building upon previous work [35,61–64], we expressed *Bma-gar-3* in the *C. elegans* body wall muscle, pharynx, and sensory neurons. Different phenotypic endpoints were optimized to measure receptor activity across these parasitized strains. Simple plate-based assays allowed for the deorphanization of *Bma*-GAR-3 expressed in the body wall and sensory neurons. We focused primarily on pharyngeal expression, given amenability to a range of visual, electrophysiological, and imaging assays. While visual observations of pharyngeal pumping and electropharyngeogram recordings provided different measurements of *Bma*-GAR-3 perturbation, these assays are ultimately low in throughput. To enable more facile and higher throughput screening of pharynx-expressed receptors, we deployed a microtiter plate imaging assay that measures the accumulation of lipophilic dye as a reporter of pharyngeal pumping. We show that this feeding assay can be used to detect activation of *Bma*-GAR-3 within the pharynx.

The suitability of these approaches for a given parasite GPCR will depend on the complement of related receptors and endogenous ligands that signal in targeted tissues. We show that transgenic strains can be created in various genetic knockout backgrounds to help mitigate some of these concerns. While expression in scalable single-cell systems will remain an important objective for high-throughput screening (HTS) against GPCR targets, functional parasite receptor assays in a more native nematode cell and physiological environment can provide important baseline pharmacological data and transgenic whole-organism assays can conceivably be adapted for high-throughput screening and anthelmintic discovery.

## Methods

### Protocol and data availability

All primary data (phylogenetic, qPCR, phenotypic) and pipelines for statistical analysis and data visualization are available at https://github.com/zamanianlab/Bm-GAR-ms.

### Parasites and chemical reagents

*Brugia malayi* and *Brugia pahangi* adults extracted from the *Meriones unguiculatus* infection system (NIH/NIAID Filariasis Research Reagent Resource Center) [102] were maintained in daily changes of RPMI 1640 with L-glutamine (Sigma-Aldrich) supplemented with FBS (10% v/v, Fisher Scientific) and penicillin-streptomycin (0.1 mg/mL, Gibco) at 37°C with 5% CO_2_ unless otherwise specified. *Brugia* microfilariae isolated from the same system were maintained in RPMI 1640 with L-glutamine (Sigma-Aldrich) supplemented and penicillin-streptomycin (0.1 mg/mL, Gibco) at 37°C with 5% CO_2_ unless otherwise specified.

Chemicals used in assays include serotonin (Fisher Scientific cat#AAB2126306), arecoline (Fisher Scientific cat# AC250130050), atropine (Santa Cruz Biotechnology cat# sc-252392), nicotine (Santa Cruz Biotechnology cat# sc-482740), levamisole (VWR cat #TCL0231-1G), carbachol (Santa Cruz Biotechnology cat# sc-202092), acetylcholine (Fisher Scientific cat# AC159170050), oxotremorine-M (Santa Cruz Biotechnology cat# sc-203656), aldicarb (Santa Cruz Biotechnology cat# sc-254939).

### Phylogenetics

Putative parasite GARs were identified using a reciprocal BLASTp [103] approach using known *C. elegans* GARs. This initial list of GARs was expanded with homology-based searches against the *C. elegans* predicted proteome and a broader list of *C. elegans* biogenic amine receptors was used to carry out blastp searches against the predicted proteomes of *B. malayi, Ancylostoma caninum, Ascaris suum, Haemonchus contortus, Strongyloides ratti,* and *Trichuris muris* (WormBase ParaSite v16 [104]). Filtered hits (percent identity > 30%, E-value < 10^-4^, and percent coverage > 40%) that survived a reciprocal blastp search against *C. elegans* were retained. This list was combined with human muscarinic receptors for phylogenetic inference and annotation. Receptors were aligned with MAFFT [105], trimmed with trimAl [106], and phylogenetic trees were inferred with IQ-TREE [107]. Trees were visualized and annotated with ggtree [108].

### Adult parasite assays

Multivariate phenotyping of adult parasites was performed as described [67,109]. After receipt from the FR3, adult male and female *B. pahangi* parasites were manually sorted into 24-well plates filled with 750 μL of complete media (RPMI 1640 + 10% FBS + pen/strep) per well. Parasites were incubated overnight, after which individual parasites were transferred to new plates with 750 μL incomplete media. 100X compound stocks were made fresh daily in H2O.

Plates were recorded for 15 sec, compound was added, and plates were immediately recorded again. Recordings were taken 1 hr post-treatment and 24 hr post-treatment, prior to transferring parasites to a new pre-loaded drug plate. Final recordings were taken 48 hr post-treatment. Three biological replicates from separate batches of parasite infection cohorts were assayed with four worms per treatment. Videos were analyzed with the optical flow (motility) module of wrmXpress [68].

Conditioned media from female worms from the initial overnight incubation in complete media and 24/48 hr treatment plates were transferred to 1.5 mL tubes and centrifuged for 5 min at 10,000 *x g* to pellet progeny. 500 μL of the supernatant was removed, and the remainder was stored at 4°C for >48 hr. To quantify progeny, 50 μL aliquots were transferred to wells of a 96-well plate (Greiner), and each well was imaged with transmitted light at 2X with an ImageXpress Nano (Molecular Devices). Images of progeny were segmented as previously described [67], and segmented pixels were counted to infer output of progeny using a previously generated model [67].

### Microfilariae assays

*Brugia malayi* microfilariae motility and cell toxicity assays were performed as described [67,110]. Briefly, mf are purified using a PD-10 Desalting Column (VWR, cat# 95017-001) into culture media and titered to a concentration of 10 mf/μL. 100 μL containing 1,000 mf were added to each well of a 96-well plate. Heat killed controls were incubating for 1 hr at 60°C before being added to wells. Serial dilutions of 100 mM stock of each drug was made fresh in water. Plates were imaged immediately after treatment (0 hr) and 24 and 48 hr after treatment. At least two replicates with high technical replication were run for motility assays. CellTox (Promega cat# G8742) staining was performed at 48 hr following the kit protocol with half the recommended concentration of CellTox. Wells were imaged using an ImageXpress Nano (Molecular Devices). Three replicate dose-response plates were assayed using CellTox assays. Videos were analyzed with the motility and segmentation modules of wrmXpress [68] and output data was analyzed using the R statistical software.

### qPCR

Parasites were flash frozen in liquid N2 in 1.5 mL tubes and stored at −80°C in TRIzol LS (Thermo Scientific cat# 10296028) in batches of 500,000 (mf), 300-500 (L3), or 3 (adults). Three independent samples originating from different batches of animal infections were collected for each stage (mf, L3, adult male, and adult female). Freeze-thawed samples were homogenized by compact bead mill (TissueLyser LT, Invitrogen) and RNA was extracted using the Zymo Direct-zol RNA Miniprep kit (cat# R2050). RNA integrity and concentration were assessed via NanoDrop, and cDNA was generated with the SuperScript III kit (ThermoFisher cat# 18080051) using equal amounts of oligo(dT) and random hexamer primers for first-strand synthesis. qPCR primers for *Bma-gar-2* (F: 5’-TAATACGACTCACTATAGGGCGACGTACTTCCTCCGATGT-3’, R: 5’-TAATACGACTCACTATAGGGCCGCTCATCGTATTCCATTT-3’) and *Bma-gar-3* (F: 5’-TTTGGCCACCATGGATTATT-3’, R: 5’-TGTATAACGCAACGGTCAGG-3’) were designed with Primer3 [111] and optimized to quantify expression levels from cDNA. GAPDH primers [112] were used as a control. 20 μL qPCR reactions were carried out using 2x PowerUp SYBR Green Master Mix (Fisher Scientific cat# A25776), 800 nM primers, and 10 ng of cDNA as input. Reactions were run in triplicate on a StepOnePlus real-time PCR system with the following program: 2 min for 50°C, 95°C for 5 min, 40 cycles of: 95°C for 15 sec, 55°C for 15 sec, and 72°C for 1 min. C_T_ values were calculated with the system’s automatic threshold.

### RNAscope *in situ* hybridization

Adult *B. malayi* females were cultured overnight and separated in 10 cm petri dishes, dipped in 70% ethanol, and spatially embedded in 1% bacto-agar. Blocks of bacto-agar were dehydrated for 5 minutes in 25%, 50%, 70% ethanol sequentially and stored in 100% ethanol overnight. The bacto-agar embedded tissue was processed into a paraffin block and agar was trimmed from the anterior tip of the agar block before embedding for cross-sections. Blocks were sectioned at 8 μm and sections were arranged sequentially on slides, keeping all sections.

Hybridization probes used for RNAscope (ACD Bio) include DapB (dihydrodipicolinate reductase, *Bacillus subtilis)* as a negative control and two highly-expressed *B. malayi* genes as positive controls: *Bma-Bm4733 (B. malayi* β-tubulin targeting 2-1031 of XM_001896580.2 (20ZZ)) and *Bma-cdc-42 (B. malayi* putative GTP-binding protein targeting 2-550 of XM_001899971.1 (11ZZ)). Our *Bma-*GAR-3 target probe targets 400-1501 of XM_043081323.1 (20ZZ). The standard RNAscope™ 2.5 HD Assay-RED kit (ACD Bio) protocol was followed with an adjusted length of 8 min for retrieval using a steamer and 45 min for amp 5. Imaging was done in bright field with 40x-Nikon Plan Apo on a Nikon Eclipse80i. Presence of red punctate staining, indicating target gene expression, was quantified by manual annotation by two independent observers. Annotation and alignment were carried out with Fiji [113] using TrakEM2 [114].

### Cloning and transgenegic *C. elegans* strains

The open-reading frame (ORF) for the longest predicted isoform of *Bma-gar-3* (isoform a, WormBase ParaSite version 17) was selected for study based on manual assessment and alignment with homologous GARs. This isoform is supported by long-read RNA sequencing data [115] that extends the model upstream of the 5’ end of isoform b. The ORF was synthesized (GenScript) and cloned into pPD133.48, L4663 (a gift from Andrew Fire, Addgene plasmid #1665) using BamHI/KpnI sites to create pMZ0005 *(myo3p::Bma-gar-3::unc-54-3’UTR).* pMZ0012 *(sra-6p::Bma-gar-3::unc-54-3’UTR)* was created by subcloning *Bma-gar-3* into *sra-6p::ChR2*YFP* (a gift from Shawn Lockery [116]) using BamHI/EcoRI sites. pMZ0018 *(myo-2p::Bma-gar-3::unc-54-3’UTR)* was created by amplifying *myo-2p* from pPD96.48, L2531 (a gift from Andrew Fire, Addgene plasmid #1607) with 5’ XbaI and 3’ BamHI sites and using this amplicon to replace *myo-3p* in pMZ0012.

Genotypes generated and used in this study include: ZAM7 *(gar-3(gk305)* V, *maz7Ex[sra-6p::Bma-gar-3::unc-54* 3’UTR; *sra-6p::GCaMP3; unc-122p::GFP]),* ZAM10 *(gar-3(gk305)* V, *maz10Ex[myo-3p::Bma-gar-3::unc-54* 3’UTR; *myo-2p::GFP]),* and ZAM11 *(gar-3(gk305)* V, *maz11Ex[myo-2p::Bma-gar-3::unc-54* 3’UTR; *myo-2p::GFP])* were created as described [117] by injecting pMZ0012 (75 ng/μL), pMZ0005 (30 ng/μL), and pMZ0018 (30 ng/μL) into *gar-3(gk305),* respectively, along with fluorescent markers *(unc-122p::GFP* or *myo-2p::GFP)* and empty vector (pPD95.75) to create a final concentration of 100 ng/μL. ZAM19 ([*myo-2p::GFP*]) and ZAM20 *(gar-3(gk305)* V *[myo-2p::GFP])* were created by injecting 10 ng/μL of a fluorescent marker *(myo-2p::GFP)* and empty vector (pPD95.75) to create a final concentration of 100 ng/μL. *VC657(gar-3(gk305)* V) was sourced from the *C. elegans* Gene Knockout Consortium [118]. Lines were maintained at 20°C on NGM plates seeded with *E. coli* strain OP50 and routinely picked to fresh plates at the L4 stage.

### Ring aversion assay

Bleach synchronized adult worms were rinsed three times in M9 and pipetted onto unseeded 6 cm plates. To assemble the assay plate, a copper ring (PlumbMaster cat# 17668) was soaked in a test compound with fast green dye as a visual marker and placed in the center of the plate. 1 μL of 1:1000 diluted diacetyl (attractant) was combined with 1 μL fast green and added outside of the ring. Plates were used immediately after assembly. The copper ring was removed after 1 minute to allow soaking of the test compound into the agar. Individual worms were picked without bacteria to the center of the compound ring on assay plates and monitored for reversals in response to test compounds. Three biological replicates from three independent bleaches were run using four worms per treatment per strain. Observations were made until either the worm crossed the compound ring or six attempts to cross the compound ring occurred without the worm crossing. Water was used as a negative control and 4 M D-fructose was used as a positive control.

### Aldicarb paralysis assay

1 mM aldicarb plates were made by diluting 100 mM aldicarb (in 70% ethanol) in NGM. Plates were stored at 4°C and used within 30 days. Thirty L4s from each strain were picked to OP50 seeded NGM plates and cultured at 20°C overnight until they reached the young adult stage. Aldicarb plates were acclimated to room temperature on the morning of the assay. Four copper rings (PlumbMaster cat# 17668) were dipped in 70% ethanol and briefly flamed before placement onto a single aldicarb plate to form quadrants. 10 μL of OP50 from overnight liquid culture was spotted into the center of each of the copper rings and allowed to dry for 30 min. A minimum of 10 worms per strain were picked to the center of a quadrant, allowing the testing of four strains per plate. Four biological replicates were carried out across strains. After 30 min and every 10 min thereafter, each worm was tapped 3 times on the head and tail, paralyzed worms were removed, and remaining worms were tallied. This process continued until the 2 hr mark.

### Visual pharyngeal pumping assay

For L1 stage assays, gravid worms were bleached and embryos were hatched on unseeded 10 cm NGM plates overnight (2,000 embryos per plate). L1s were washed from plates into 1.5 mL tubes and treated with drug for 20 min. 500 μL of the treated worm mixture was then transferred to an unseeded 10 cm NGM plate and allowed to dry for 5 min. Individual transgene (+) worms were then picked to seeded 6 cm NGM assay plates and given 10 min to acclimate. Adult assays required minor modifications of the L1 protocol. Young adults were developed on seeded 10 cm NGM plates, drug-treated worms were transferred to 10 cm plates with no drying period, and transgene (+) worms were picked to unseeded 6 cm NGM assay plates. For optimization and cholinergic assays, an average of 8 worms were visually phenotyped for each strain-condition combination. Pumps were measured by the motion of the terminal bulb grinder over a 30 sec (L1s) or 10 sec (adults) interval using a Zeiss Axio Vert.A1 at 10x with DIC optics.

### Electropharyngeogram recordings

Electropharyngeograms (EPGs) were recorded with the ScreenChip system (InVivo Biosystems). Briefly, worms were bleach synchronized and embryos were hatched and developed on seeded 10 cm NGM plates. Young adult worms were washed 3 times with M9 in 1.5 mL tubes. Worms were treated for 20 min before loading them into the ScreenChip40 microfluidic chamber. Each worm was vacuumed into the channel, acclimated for 30 sec, and recorded for 2 min using the NemaAcquire 2.1 software. Worms from three biological replicates representing independent bleaches were assayed for optimization (minimum of 15 worms per condition), and one replicate was assayed for the cholinergic panel (minimum of 5 worms per condition). Analysis was done using NemaAnalysis 0.2 with default settings found under customize analysis. Filtering of data was done by modifying “parameters left” for signal to noise ratio (SNR), E and R highpass cutoffs, and minimum absolute threshold followed by visually ensuring all pumps were identified and background noise was ignored.

### Image-based feeding assay

Bleach synchronized larval stage worms (L1) were aliquoted into 96-well plates at a titer of 50 L1 per well along with *E. coli* HB101 bacterial food (2.5 mg/mL final concentration). Worms were incubated for 48 hr at 20°C, shaking at 180 RPM until reaching adult stage. For the initial screen, adult stage worms were drug treated for 20 min followed by addition of 180 ng/mL BODIPY 558/568 (Thermo Scientific cat#D3835) combined with HB101 and again incubated for 20 min. For optimization assays, 45 ng/mL (1x) or 90 ng/mL (2x) BODIPY 558/568 combined with M9 or HB101 was added to the plate with the same incubation time. Plates were washed 3 times with M9 and paralyzed with 50 mM sodium azide. Images of worms in transmitted light, GFP, and TxRed were taken on an ImageXpress Nano (Molecular Devices) at 2x. Images were analyzed with the feeding module of wrmXpress, which incorporates a custom CellProfiler model using the Worm Toolbox plugin [119,120]. Segmented worms were computationally straightened and GFP and TxRed were quantified. Transgenic worms were identified using a robust cutoff of GFP quantification such that only worms expressing *Bma-gar-3* were analyzed (**Sup Figure 2 A-B**). Non-worm objects and contaminating fluorescence were further filtered with size thresholds and outlier pruning (TxRed FLU z-score > 3). Total fluorescent dye uptake for GFP(+) worms was quantified using the StdIntensity parameter.

## Acknowledgements

The authors thank the University of Wisconsin Translational Research Initiatives in Pathology laboratory (TRIP), supported by the UW Department of Pathology and Laboratory Medicine, UWCCC (P30 CA014520) and the Office of The Director-NIH (S10 OD023526) for use of its facilities and services. Some strains were provided by the CGC, which is funded by NIH Office of Research Infrastructure Programs (P40 OD010440). *Brugia* life cycle stages were obtained through the NIH/NIAID Filarial Research Reagent Resource Center (FR3), morphological voucher specimens are stored at the Harold W. Manter Museum at University of Nebraska, accession numbers P2021-2032. Image processing was performed using the compute resources and assistance of the UW-Madison Center For High Throughput Computing (CHTC) in the Department of Computer Sciences. The CHTC is supported by UW-Madison, the Advanced Computing Initiative, the Wisconsin Alumni Research Foundation, the Wisconsin Institutes for Discovery, and the National Science Foundation, and is an active member of the OSG Consortium, which is supported by the National Science Foundation and the U.S. Department of Energy’s Office of Science. The authors would also like to thank the members of the Zamanian laboratory for critical comments on the manuscript.

## Supplemental Material

**Supplemental Figure 1** Effects of cholinergic treatment on electrophysiological phenotypes measured using EPG recordings. Duration measures are reported in ms, amplitude = μV, and frequency = Hz.

**Supplemental Figure 2** A) Data was manually annotated by visually labeling worms in wrmXpress output images as GFP+ or GFP- (transgenic strain populations created with extrachromosomal arrays contain both GFP+ and GFP-populations indication expression of transgenes). Each GFP-related measure was plotted to determine a metric for filtering data for only GFP+ (transgene expressing) worms. B) StdIntensity, used to filter GFP+ worms from mixed populations, shows a clear separation between GFP+/- populations at a threshold of .0015.

