## Supplementary figures and images for "Pharmacological profiling of a *Brugia malayi* muscarinic acetylcholine receptor as a putative antiparasitic target"

### Supplementary Figure 1

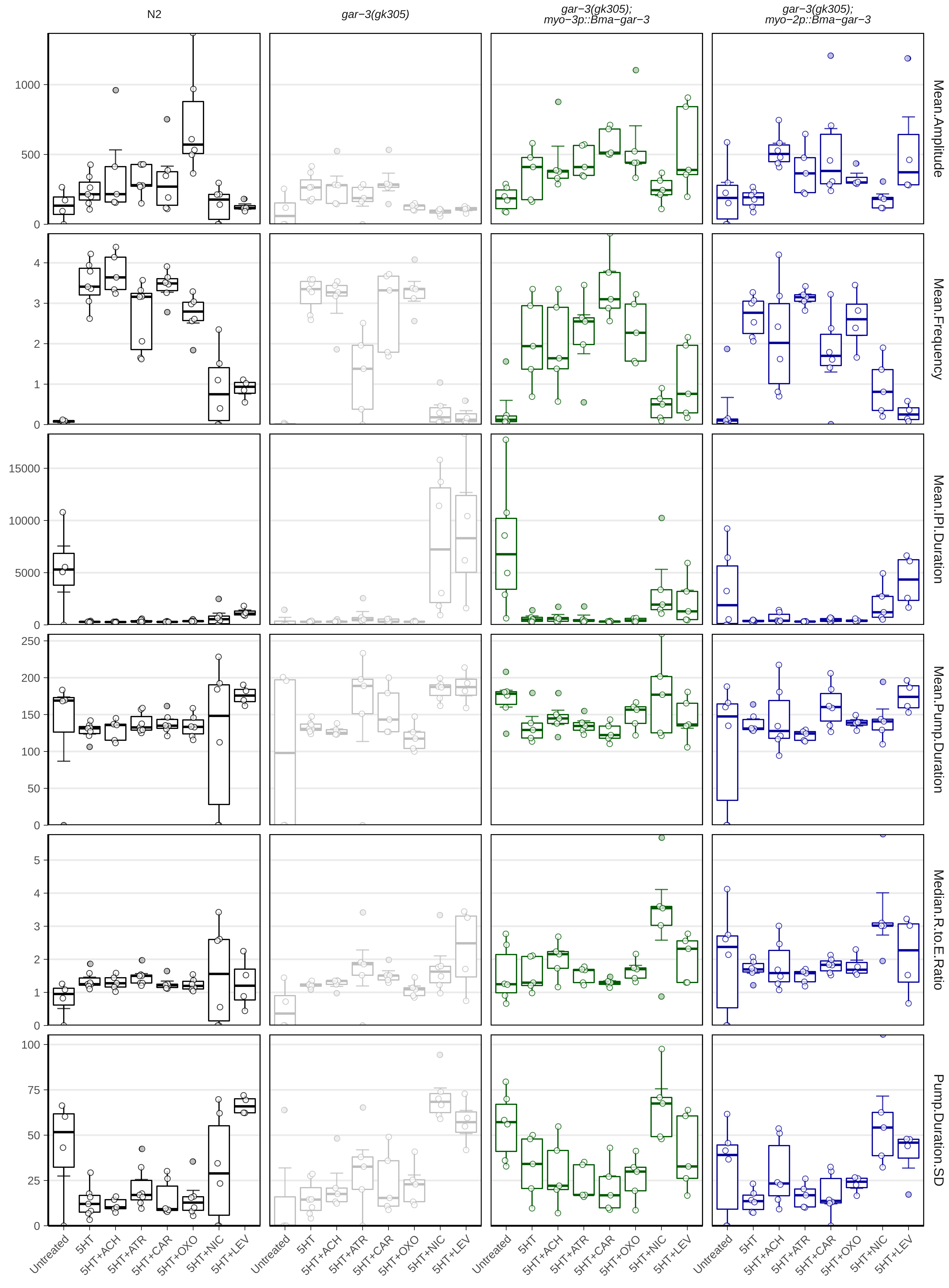

### Supplementary Figure 2

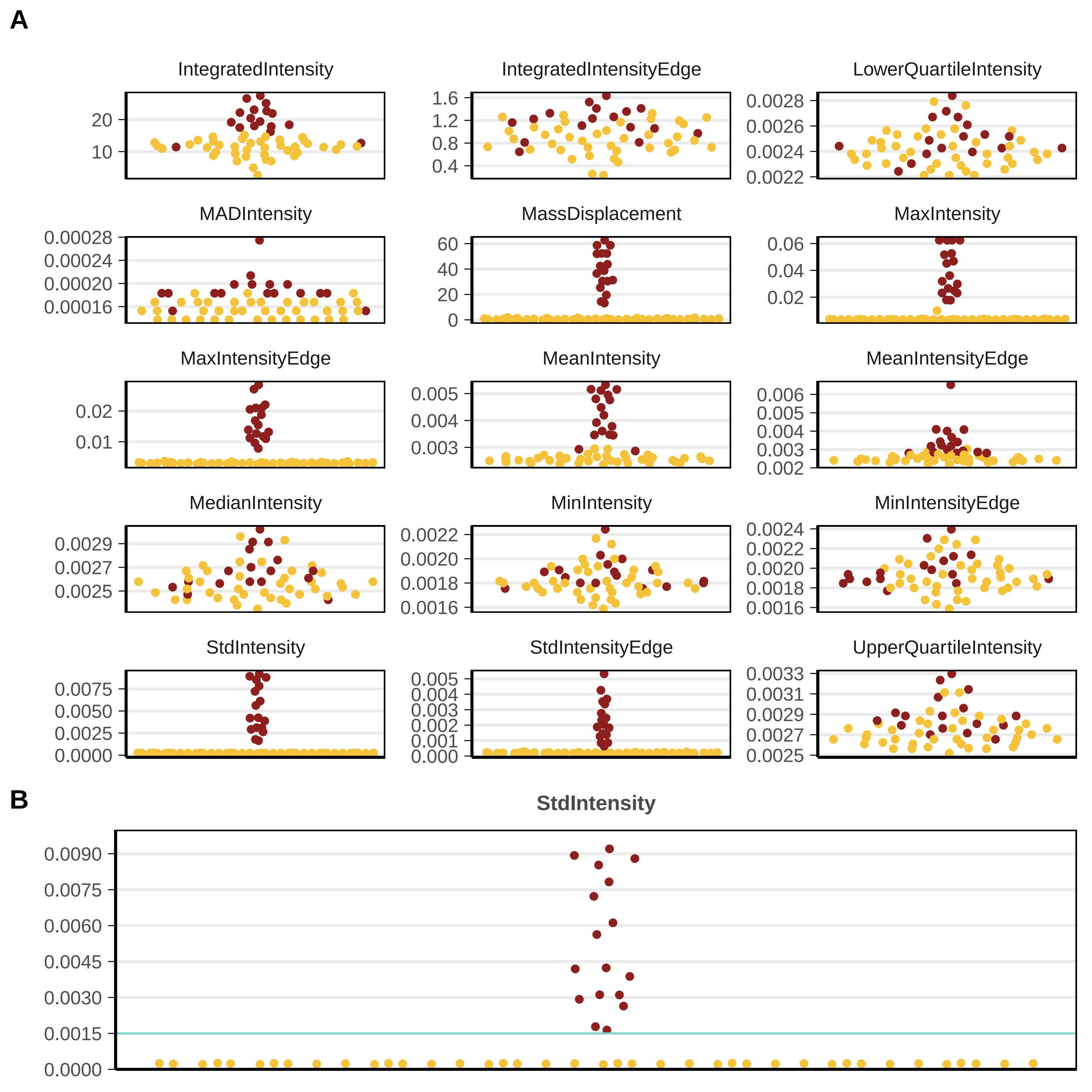
